# CDK inhibitors reduce cell proliferation and reverse hypoxia-induced metastasis of neuroblastoma tumours in a chick embryo model

**DOI:** 10.1101/405639

**Authors:** Rasha R. Swadi, Keerthika Sampat, Anne Herrmann, Paul D. Losty, Violaine See, Diana J. Moss

## Abstract

Neuroblastoma is a paediatric cancer with a poor prognosis. This is in part due to the widespread metastasis at time of presentation, which is refractory to current treatment modalities. New therapeutic agents that can control not only tumour growth but also metastasis are urgently needed.

One current therapeutic option used in the clinic is differentiation therapy with retinoic acid, where the terminal differentiation of the neuroblastoma cells reduces tumour growth in the primary tumour as well as at metastatic sites. However, retinoic acid only works in a subset of patients.

We investigated the potential of CDK inhibitors on neuroblastoma cell differentiation, tumour progression and metastasis by utilising a 3R compliant cost effective preclinical chick embryo model. In both SK-N-AS and BE(2)C cell lines, when engrafted on the chorioallantoic membrane of chick embryos, we observed a reduction of tumour cell proliferation as well as a reduction in hypoxia preconditioning-driven metastasis by 60%. In addition, the expression of a panel of genes with known roles in metastasis, which increased upon hypoxia-preconditioning, was largely reduced by a CDK1 inhibitor. These results provide a promising alternative to currently existing therapies and might aid the development of new treatment protocols for retinoic acid-resistant patients.

## Background

Neuroblastoma is a paediatric cancer that originates due to delayed development of the neural crest cells destined to form the sympathetic nervous system or adrenal glands [1]. Most patients present with high risk metastatic disease. However patients presenting before 18 months of age have a better prognosis, despite having widespread disease. This is thought to be due in part to the eventual differentiation of tumour cells in younger patients, and has led to the development of differentiation therapy, such as retinoic acid, as a strategy for treating neuroblastoma [2, 3]. Although the inclusion of retinoic acid in treatment protocols results in enhancement in survival, many patients still relapse. In culture, cell lines with amplification of the MYCN gene typically respond to retinoic acid whilst cell lines or patients with other genetic modifications such as loss of chromosome 11q do not [4], hence the need for additional treatment options for this group.

Retinoic acid acts by modifying gene expression via complex formation with its receptor RAR and downregulation of MYC-N levels. Hence, it is most effective in inducing differentiation in tumours with high MYCN expression [5, 6]. Retinoic acid also decreases cell proliferation by increasing levels of p27(Kip1), an inhibitor of cyclin D kinase (CDK) 2, 4 and 6 [7]. Decreasing proliferation is thought to be intimately associated with the initiation of differentiation during normal development [8] and this led us to hypothesise that agents directly inhibiting CDKs may provide an additional or alternative approach to promoting differentiation of neuroblastoma cells. CDK inhibitors have previously been tested in a range of tumour types including breast cancer where Palbociclib, a CDK 4/6 inhibitor, has been approved for use in patients [9, 10]. RO-3306, a CDK1 inhibitor, has also been successfully tested in preclinical models [11, 12].

We have previously developed a chick embryo model to enable the investigation of neuroblastoma growth and its metastatic process. We demonstrated that several neuroblastoma cell lines form tumours on the chick embryo chorioallantoic membrane (CAM) [13]. We have further shown that upon hypoxia preconditioning (1% O_2_) for three days prior to implantation, cells metastasised into the chick embryo from 60% of the tumours observed [13]. This model has further been shown to be suitable for therapeutic agents testing [14]. We therefore used it to test whether CDK inhibitors, known to slow cell proliferation, may also reduce or prevent metastasis.

## Results

### CDK4/6 inhibitor promotes differentiation of BE(2)C cells while CDK1 inhibitor promotes cell death

Previous work has shown that all-trans retinoic acid (ATRA) promotes cell differentiation and reduces cell proliferation in some neuroblastoma (NB) cell lines, possibly by increased expression of p27Kip1, an inhibitor of CDK2, 4 and 6 [7]. CDK inhibitors, which are known to slow the cell cycle, may therefore promote the differentiation of neuroblastoma cells. To test this hypothesis, BE(2)C cells were treated with the CDK4/6 inhibitor Palbociclib (CDK4/6i) for 3 days, alone or in combination with ATRA to test whether the CDK4/6i might enhance the ATRA effects on neuroblastoma differentiation. In terms of morphology, while 5 μM of CDK4/6i promoted the growth of cell extensions similar to those seen with ATRA, doses of 10 μM and above promoted cell death (Figure 1A). Treatment with 5 μM CDK4/6i was enough to reduce the proliferation of BE(2)C cells by 59% essentially similar to the ATRA effects (Figure 1B). The combination of the CDK4/6i and ATRA did not further decrease cell proliferation (Figure 1C), suggesting that these 2 drugs have similar mechanisms of action. To confirm that CDK4/6i was driving BE(2)C cell differentiation, the expression of previously characterised markers KLF4, ROBO2 and STMN4, [14] was measured. In all three conditions (drugs alone or in combination), the expression of the stem cell marker KLF4 decreased whilst the expression of the differentiation markers ROBO2 and STMN4 increased compared to untreated cells. Again, CDK4/6i was no more potent than ATRA alone (Figure 1D). The expression of MYCN was not significantly altered in response to the CDK4/6i, whereas a small decrease was observed in response to ATRA (Figure 1D).

**Figure 1.**
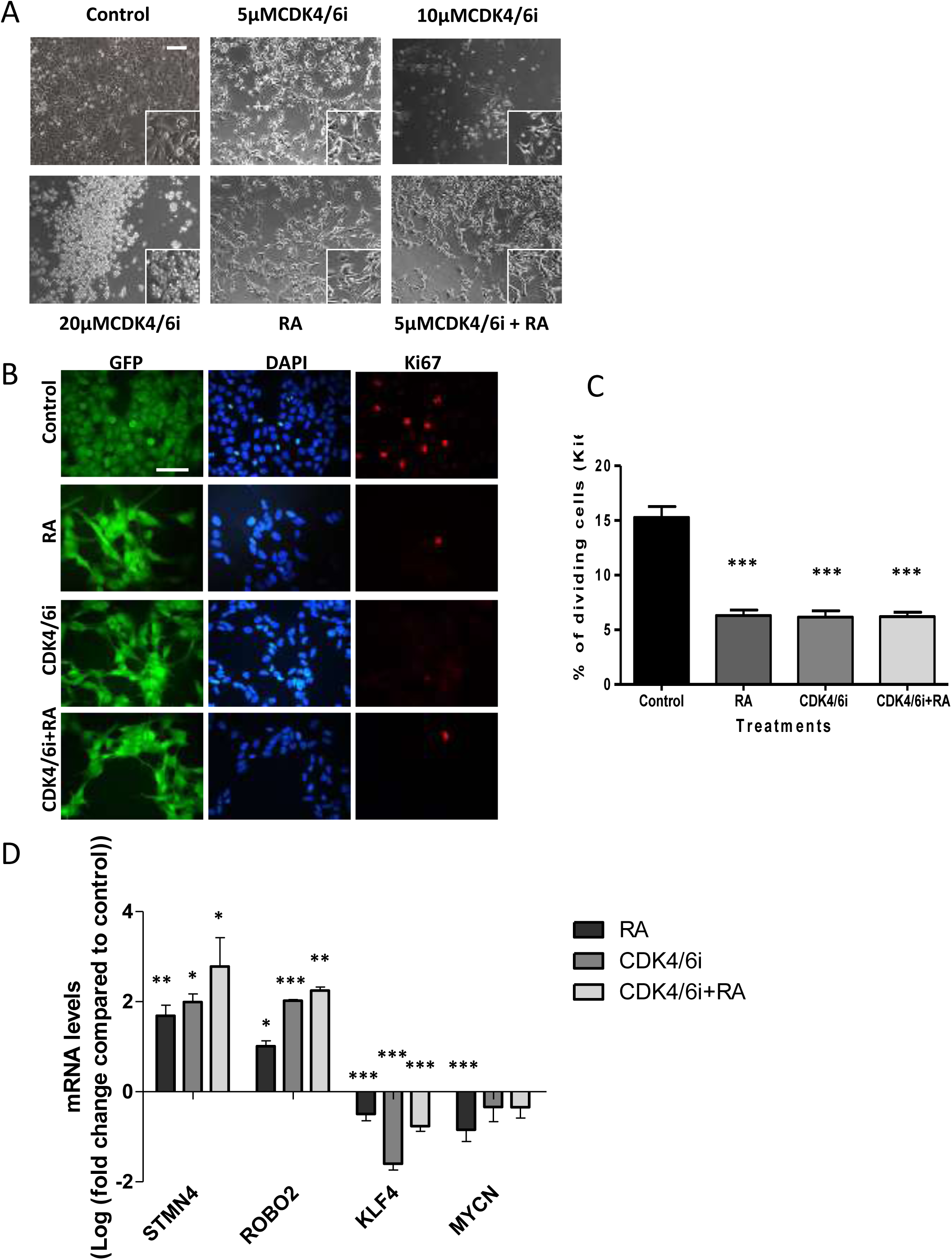
The CDK4/6i and ATRA promote differentiation of BE(2)C cells and CDK4/6i induces death at 10 and 20 μM. **A** BE(2)C cells were treated for 3 days with DMSO, 5 μm, 10 μm, 20 μm CDK4/6i, 10 μm ATRA and 5 μm CDK4/6i and 10 μm ATRA combined. 5 μm CDK4/6i, 10 μm ATRA and the combination all showed a change in morphology including cellular extensions of variable length. 10μm and 20 μm CDK4/6i showed increasing numbers of floating cells indicative of cell death. **B** GFP-expressing BE(2)C cells were grown for 3 days in 5 μm CDK4/6i, 10 μm ATRA and both combined and then stained for Ki67 (red) and DAPI (blue). **C** Quantification of Ki67-positive cells as a percentage of the total cell number. All three treatments showed a 59% reduction in cell proliferation. Each bar represents the mean ±SEM of three independent experiments (n = 3) and at least 9 fields per experiment. *** P≤0.001 compared with the control. Scale bar =100 μM. **D** Relative mRNA levels for the target genes were determined by qPCR. Cells were cultured with either 5 μM of CDK4/6i, 10 μM of ATRA or in combination for 3 days. At least three independent experiments (n = 3) were analysed for each condition and mRNA levels are displayed relative to GAPDH, UBC and HPRT1 and normalised to cells cultured for 3 days with DMSO (ATRA) or PBS (CDK4/6i). Each bar in the graph represents the normalised mean± SEM of three independent experiments. *P≤0.05, **P≤0.01 and ***P≤0.001 compared with control.

CDK4/6 acts at the G1/S boundary, whereas CDK1 acts at both the G1/S and G2/M boundaries. We therefore further tested whether a CDK1 inhibitor would have a similar effect to the CDK4/6 inhibitor. BE(2)C cells were treated for 3 days with RO-3306, a CDK1 inhibitor. Based on morphology, unlike CDK4/6i, RO-3306 promoted cell death in contrast to inhibition of proliferation and/or promotion of differentiation even at 5 μM (Figure 2A). At 1 μM there was no change in morphology and no evidence for cell death (data not shown). Twenty four hour treatment of BE(2)C cells with 5μ M RO-3306 followed by two days without drug also resulted in cell death rather than any morphological indication of differentiation (data not shown). To quantify this further, we determined cell viability after 3 days and showed that RO-3306 was indeed effective at reducing cell viability (Figure 2B). Three day treatment with 5 μM CDK1i reduced cell viability by 65% and increasing the concentration of the inhibitor did not enhance this further. Using Annexin V labelling, we demonstrated that the cell death was apoptotic (Figure 2C). We also noticed that 5 μM CDK4/6i, shown above to promote differentiation, also prompted cell death by apoptosis of some cells (Figure 2C).

**Figure 2.**
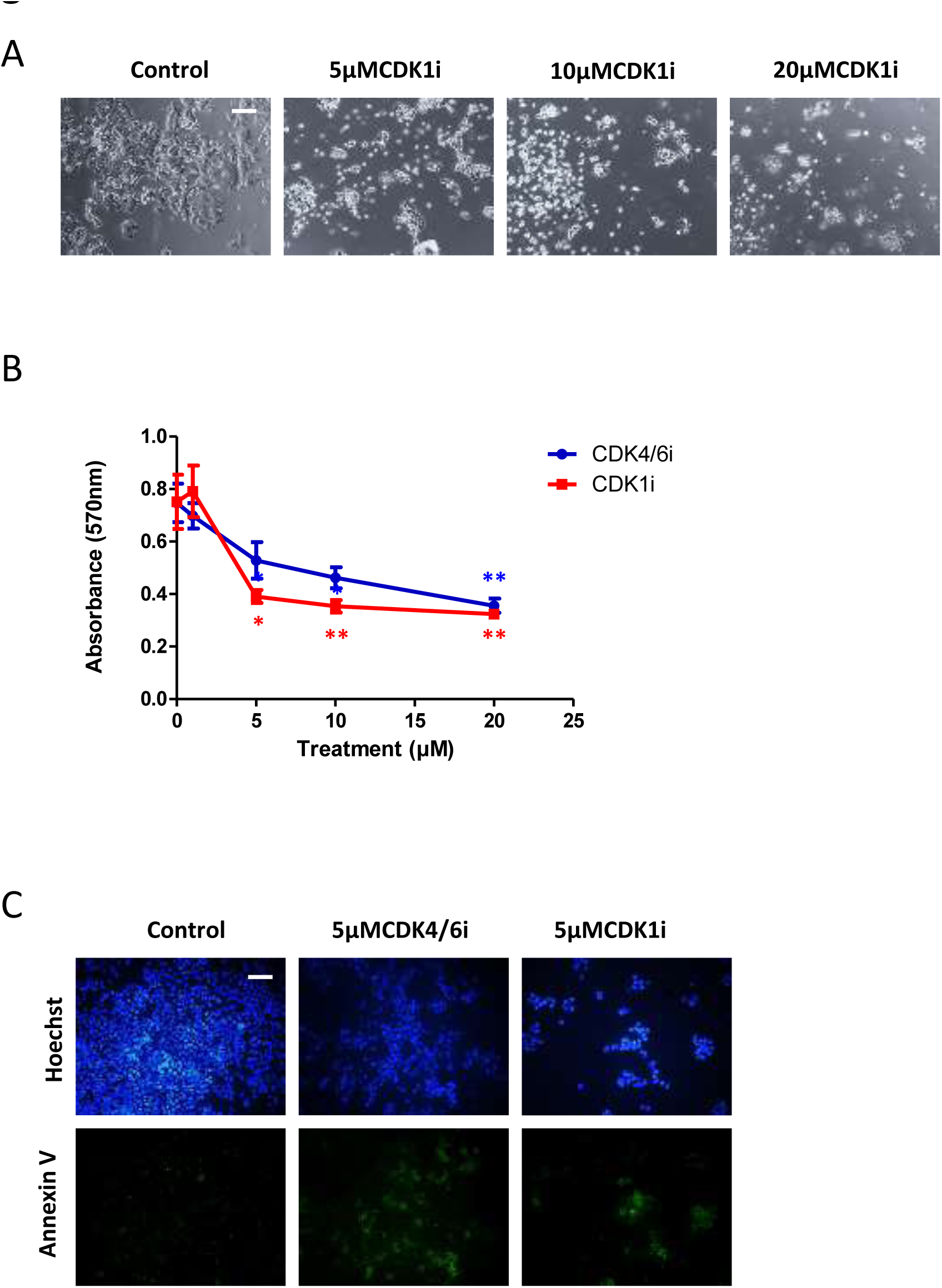
CDK1i reduces cell viability and induces apoptosis in BE(2)C cells. **A** BE(2)C cells were treated for 3 days with DMSO, 5 μm, 10 μm, 20 μm CDK1i. Increasing CDK1i concentration resulted in increasing numbers of floating cells indicative of cell death. **B** Cells were cultured for 72 h with 1, 5, 10, and 20 μM of CDK4/6i, CDK1i, or PBS (CDK4/6i) or DMSO (CDK1i) as control. MTT assay was carried out and the OD was measured at 570 nm. Data are the mean ±SEM of five independent experiments (n = 5), with 3 technical replicates for each treatment. *P≤0.05 and **P≤0.01. **C** Cells were treated for 72 h with media containing either CDK4/6i (5 μM) or CDK1i (5 μM) or DMSO as control. Cells undergoing apoptosis were visualised with Annexin V-FITC (green) and cells were identified with Hoechst 33342 (blue). Scale bar =100 μM.

### CDK4/6 and CDK1 inhibitors promote cell death of SK-N-AS cells

SK-N-AS cells express MYCN but do not have amplification of the MYCN gene. Instead they have an 11q deletion and thus represent the genetic signature of another group of patients with poor prognosis [4]. SK-N-AS cells do not differentiate or indeed respond to ATRA alone [15] so being able to target 11q-cells with an alternative approach would be very beneficial. As shown in Figure 3A the response of the SK-N-AS to both CDK4/6i and CDK1i was cell death at 5 μm, with no evidence of a change in morphology towards a differentiation phenotype at lower doses (1 μm). Cell viability was assessed after three days and again CDK1i prompted extensive cell death (58%) at 5 μM with no significant increase at higher concentrations (Figure 3B). CDK4/6i was less efficient in inducing cell death at 5 or 10 μM (Figure 3B), but there was still evidence of increased apoptosis at 5 μM (Figure 3C).

**Figure 3.**
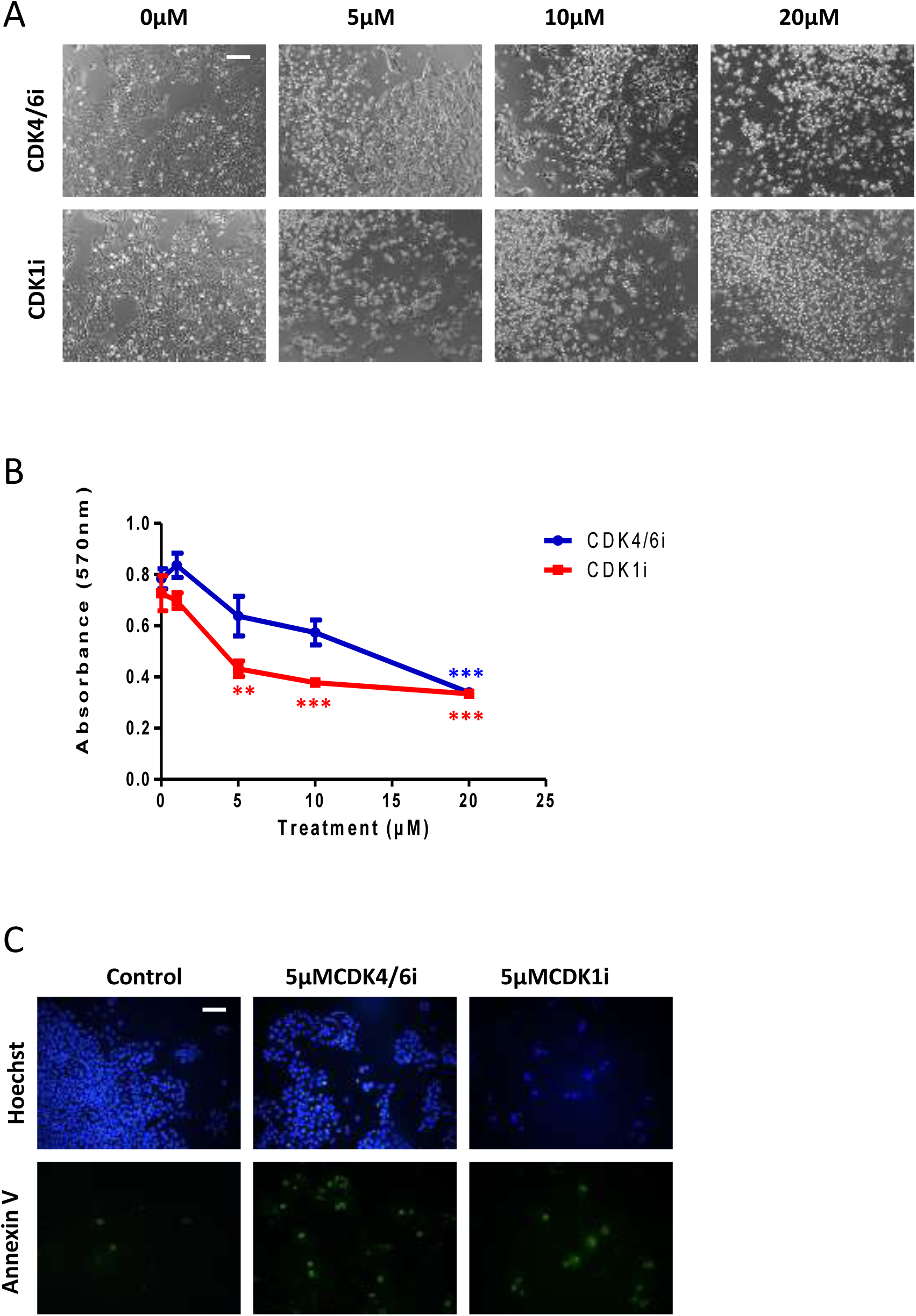
Both CDK4/6i and CDK1i induce apoptosis in SK-N-AS cells. **A** Cells were cultured for 3 days with 5 μM, 10 μM, and 20 μM CDK4/6i, CDK1i, or PBS/DMSO control. Increasing inhibitor concentration resulted in increasing numbers of floating cells indicative of cell death. **B** Cells were cultured for 72 h with 1 μM, 5 μM, 10 μM, and 20 μM of CDK4/6i, CDK1i, or PBS/DMSO control and measured at 570 nm. MTT assay was carried out and the OD was measured at 570 nm. Data are the mean ±SEM of five independent experiments (n = 5), with 3 technical replicates for each treatment.**P≤0.01 and ***P≤0.001 compared with control. Cells were treated for 72 h with media containing either CDK4/6i (5 μM) or CDK1i (5 μM) and or DMSO as control. Cells undergoing apoptosis were visualised with Annexin V-FITC (green) and cells were identified with Hoechst 33342 (blue). Scale bar =100 μM.

### CDK4/6 and CDK1 inhibitors reduce cell proliferation in both BE(2)C and SK-N-AS tumours

Cells can respond differently to drugs in a 3D *in vivo* environment compared to a 2D culture, so it was essential to test the efficacy of these drugs on tumours formed *in vivo*. We have used the chick CAM model, which not only allows robust neuroblastoma tumour formation, but also the control of their metastatic phenotype through hypoxic preconditioning [13]. We have previously shown that using four times the *in vitro* concentration of ATRA in the chick, tumours had significantly reduced number of proliferating cells, with cells also expressing differentiation markers [14]. We injected CDK inhibitors to give a concentration of 20 μM (in the egg), after tumour formation at embryonic day 11 (E11) and E13 and the treated tumours were harvested at E14. The chick embryos tolerated the doses well with no significant change in embryo survival (data not shown). Ki67-staining of BE(2)C tumour sections revealed a reduction in cell proliferation in response to both CDK4/6i and CDK1i (Figure 4A and C). CDK4/6i reduced cell proliferation of BE(2)C cells by 35%, similar to ATRA (39%) (Figure 4B). Contrary to *in vitro* data, CDK1i proved more efficient than CDK4/6i, reducing cell proliferation by almost 50%. Similar experiments were carried out with SK-N-AS cells. Since ATRA has no effect on SK-N-AS [15], only the two CDK inhibitors were tested. The reduction in proliferation was similar to that seen with BE(2)C cells (40% and 50% for CDK1i and CDK4/6i respectively; Figure 4D).

**Figure 4.**
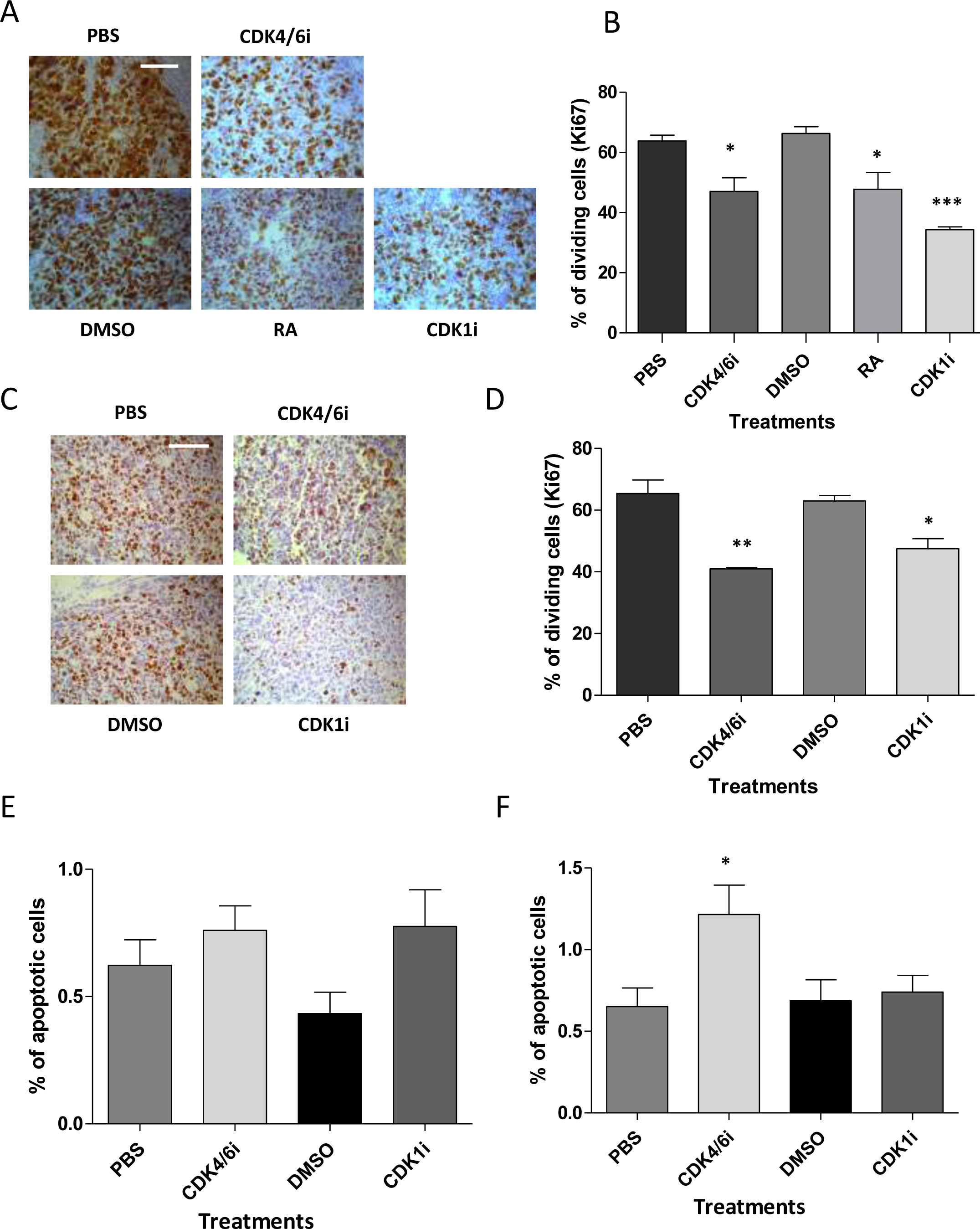
CDK4/6i and CDK1i reduce cell proliferation of BE(2)C and SK-N-AS cells within tumours. **A** GFP-labelled BE(2)C cells were implanted on the CAM of E7 chick embryos. Two injections of 0.2 ml of 9 mM ATRA, 4.5 mM CDK4/6i, 4.5 mM CDK1i, 14% DMSO, PBS, 32.5% DMSO as control respectively were made into the allantoic sac of embryos at E11 and E13. Dissected tumours were formalin-fixed and paraffin embedded and 4μm sections were stained with Ki67 (brown). ATRA, and both CDK1i and CDK4/6i reduced the number of Ki67 positive tumour cells **B** Quantification of Ki67-positive cells as a percentage of the total cell number indicates a reduction in cell proliferation for each of the three treatments. Each bar represents the mean ±SEM of three independent experiments and at least 9 fields counted per experiment, *P≤0.05 and ***P≤0.001 Scale bar is 100 μM. **C** formalin-fixed and paraffin embedded (FFPE) SKNAS sections stained with Ki67. GFP labelled SKNAS cells were implanted on the CAM of E7 chick embryos. Two injections of 0.2 ml of 4.5 mM CDK4/6i, 4.5 mM CDK1i, or PBS, 32.5% DMSO as control were into the allantoic sac of embryos at E11 and E13. 4 μm FFPE sections were stained with Ki67 (brown). Both CDK1i and CDK4/6i reduced the number of Ki67 positive tumour cells. **D** Quantification of Ki67-positive cells as a percentage of the total cell number indicates a reduction in cell proliferation for both treatments. Each bar represents the mean ±SEM of three independent experiments (n = 3) and at least 9 fields counted per experiment, *P≤0.05 and **P≤0.01. Scale bar =100 μM. **E** Quantification of TUNEL-positive cells as a percentage of the total cell number of BE(2)C cells within the section, the data is from three tumours and 9 fields per tumour and is shown here as mean ± SEM (n = 3). **F** Quantification of TUNEL-positive cells as a percentage of the total cell number SK-N-AS cells within the section, the data is from three tumours and 9 fields per tumour and is shown here as mean ± SEM (n = 3).).*P ≤0.05 compared with control.

Since, both inhibitors had a marked effect on cell survival and triggered apoptosis in cultured cells, we used TUNEL staining to evaluate the amount of apoptosis in the tumours. The number of cells dying by apoptosis was very small in control tumours (<2%). This result was consistent with the number of apoptoticcells observed by histology (Supplementary figure). None of the inhibitors triggered a significant increase in either BE(2)C or SK-N-AS tumours (Figure 4 E,F). Thus, unlike in *in vitro* cell culture, the main effect of the CDK inhibitors was a reduction in cell proliferation, rather than an increase in apoptosis

### CDK inhibitors reduce hypoxia/DMOG-induced metastasis

We next investigated the effects of the CDK inhibitors on neuroblastoma metastasis. We have previously shown that preconditioning neuroblastoma cell lines for three days in 1% O_2_ or pretreatment for 24 h with the hypoxia mimetic drug DMOG promotes metastasis of cells from the tumour formed on the CAM into the embryo [13]. Prior to the *in vivo* investigation, we initially tested whether ATRA could reverse the effect of DMOG on the BE(2)C cells proliferation *in vitro*. As shown in Figure 5A and B, DMOG increased cell proliferation by 45% *in vitro* and ATRA reversed this increase, by reducing proliferation to control levels, although not to the level of ATRA treatment on non-DMOG treated cells. The effect of the CDK inhibitors on the viability of control and DMOG pre-treated cells was also tested. Both inhibitors reduced the viability of cells (BE(2)C and SK-N-AS) irrespective of the DMOG treatment (Figure 5C & D).

**Figure 5.**
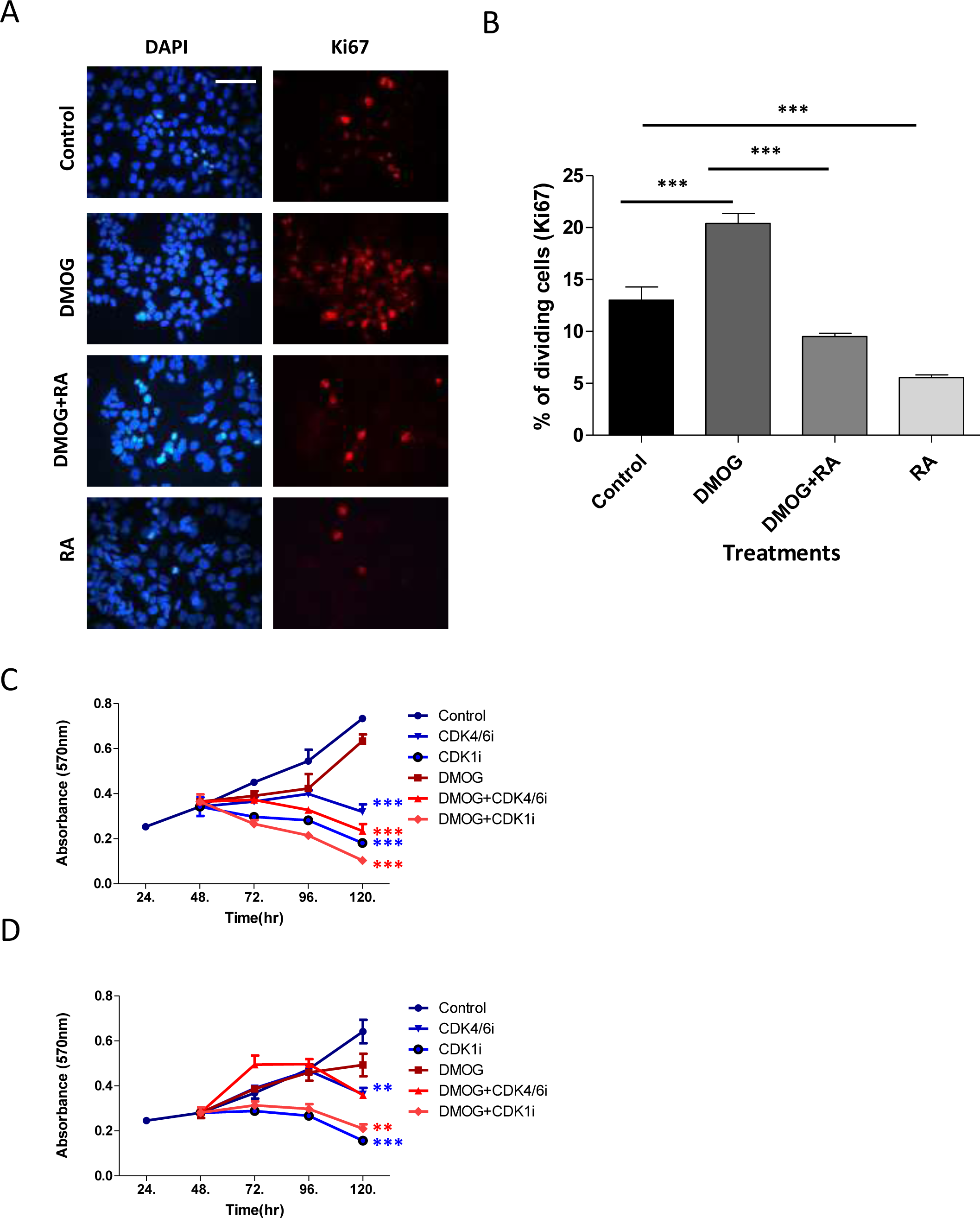
The effect of ATRA, CDK4/6 and CDK1 inhibitors on DMOG-treated BE(2)C and SK-N-AS cells. **A** BE(2)C cells were pretreated with DMOG for 24hr followed by 3 days with or without 10 μM ATRA. For comparison cells without DMOG were also treated with ATRA. Cells were then stained with Ki67 (red) and DAPI stained (blue) Scale bar =100 μM. **B** Quantification of percentage of Ki67 positive cells indicates a reduction in cell proliferation following treatment with 10 μM of ATRA in both DMOG-treated and control BE(2)C cells. Each bar represents the mean ±SEM of three independent experiments (n = 3) and at least 9 fields per experiment. ***p≤0.001 compared with the control. **C** BE(2)C cells were grown +/- DMOG for 24 h and then treated with CDK4/6i, CDK1i or medium with now drug for 3 days. The number of viable cells was assessed using the MTT assay.CDK4/6i and CDK1i both reduced the cell number irrespective of whether the cells were precondition in DMOG. Displayed is the mean ±SEM of at least three independent experiments (n = 3), with 3 technical replicates in each treatment.*P≤0.05 and **P≤0.01 compared with the control. **D** Same as for **C** but with SK-N-AS cells.

We further tested the CDK inhibitors and ATRA, for their ability to block the metastatic process *in vivo*. We have previously shown that by pre-treating cells with DMOG for 24 h or growing cells in 1% O_2_ for 3 days prior to implanting the cells on the CAM, triggered metastasis in 60% of the cases, often in the liver and gut [13]. Such a metastatic phenotype was never observed with tumours formed from cells grown in normoxia. Before testing the effect of the drugs we first needed to determine the onset time of metastatic dissemination by dissecting the embryos at E9 – E13.

Tumour formation could first be detected at E11 for BE(2)C cells and E10 for SK-N-AS cells, yet in most cases metastatic cells were only observed after E12. We therefore changed the treatment regime from injections at E11 and E13 to E10 and E12 to ensure that the inhibitors were introduced before to the onset of metastasis.

Tumours were formed from SK-N-AS cells that had been pre-treated with DMOG for 24 h. Approximately 80% of tumour cells were Ki67 positive in the DMOG tumours, a proportion similar in tumours formed by cells that had not been exposed to DMOG (Figure 4D and & 6A). Both CDK inhibitors reduced cell proliferation in the DMOG tumours by 24-39% for cdk4/6i and cdk1 I respectively (Figure 6A).

**Figure 6.**
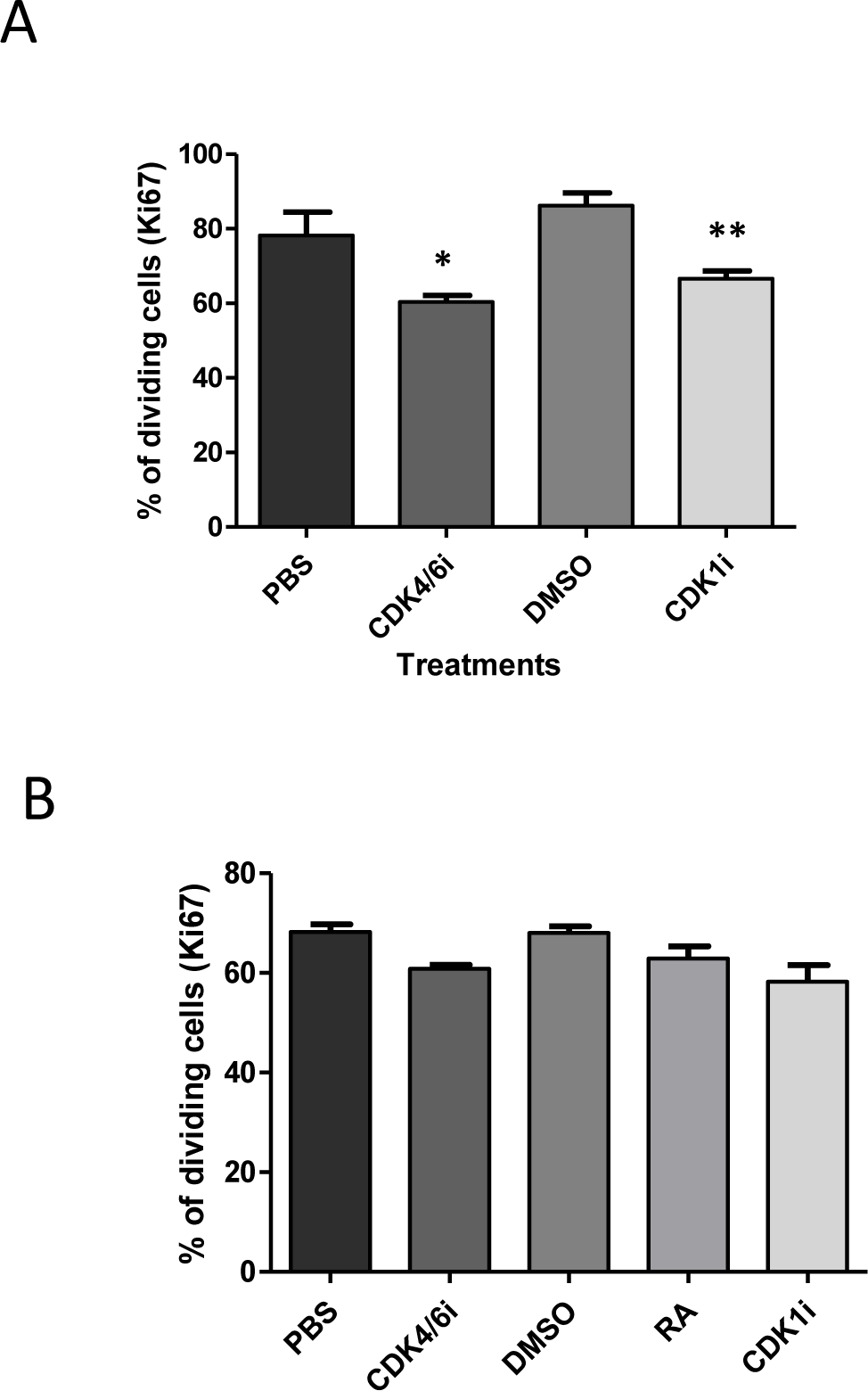
The effect of CDK4/6i and CDK1i on cell proliferation of hypoxia-preconditioned SK-N-AS and BE(2)C cells in tumours. **A** SK-N-AS were pre-treated with DMOG for 24 hr prior to implantation at E7 on the CAM of chick embryos. Two injections of 200 μl of 4.5 mM CDK4/6i or 4.5mM CDK1i, or PBS, 32.5% DMSO as control respectively were made into the allantoic sac of embryos at E10 and E12. Tumours were sectioned and stained for Ki67. The percentage of Ki67 positive cells is shown and each bar represents the mean ±SEM of three independent experiments (n = 3) and at least 9 fields per experiment.*P≤0.05 and **P≤0.01 compared with the control. **B** Quantification of Ki67-positive cells out of the total cell number BE(2)C forming-tumours. BE(2)C cells were cultured in 1% O2 for 3 days prior to implantation at E7 on the CAM of chick embryos. Two injections of 9 mM RA,4.5 mM CDK4/6i, 4.5 mM CDK1i, or 6.6% DMSO, PBS, 32.5% DMSO as controls respectively were made into the allantoic sac of embryos at E10 and E12. Tumours were sectioned and stained for Ki67. The percentage of Ki67 positive cells is shown and each bar represents the mean ±SEM of three independent experiments (n = 3) and at least 9 fields per experiment.

BE(2)C cells were overall less efficient at forming tumours on the CAM than SK-N-AS cells (70% vs 90% for SK-N-AS) and DMOG pre-treatment affected the efficiency of tumour formation for these cells. We therefore replaced the DMOG pre-treatment by hypoxia preconditioning (1% O2 for 3 days). Approximately 65% of tumour formed by cells subjected to hypoxia were Ki67 positive compared to 70% for tumours formed by cells grown in normoxia (Figure 4B and 6B). Upon treatment with the cdk inhibitors, a small decrease in proliferation was observed although none was significant (Figure 6B). Thus a similar but smaller effect was observed with these agents compared to tumours formed from cells grown in normoxia (Figure 4C).

DMOG-pretreated SK-N-AS cells efficiently formed tumours and more than 95% of embryos with tumours on the CAM, also had cells or small clumps of cells disseminated in the chick organs at E14 (detected by fluorescence upon dissection). The liver and gut were typical locations for metastatic cells, with occasional occurrence in the meninges (Figure 7A). We initially observed that DMSO alone had an impact on metastasis occurrence. DMSO above 100 μl in total is reported to compromise chick survival [16] and although chick survival was not affected in our hands with a final volume of 130 μl, it did reduce the ability of tumour cells to metastasise. We reduced the number of injections from two to one and with a single injection of CDK1i or DMSO at E10, the effects of DMSO were comparable to PBS controls and significantly different to the CDK1i (Figure 7C; 80% metastasis with DMSO alone versus 33% with CDK1i). CDK4/6i was dissolved in PBS, but for comparison, we also gave only one injection at E10. In this case the metastasis frequency was reduced from 92% to 44% (Figure 7C).

**Figure 7.**
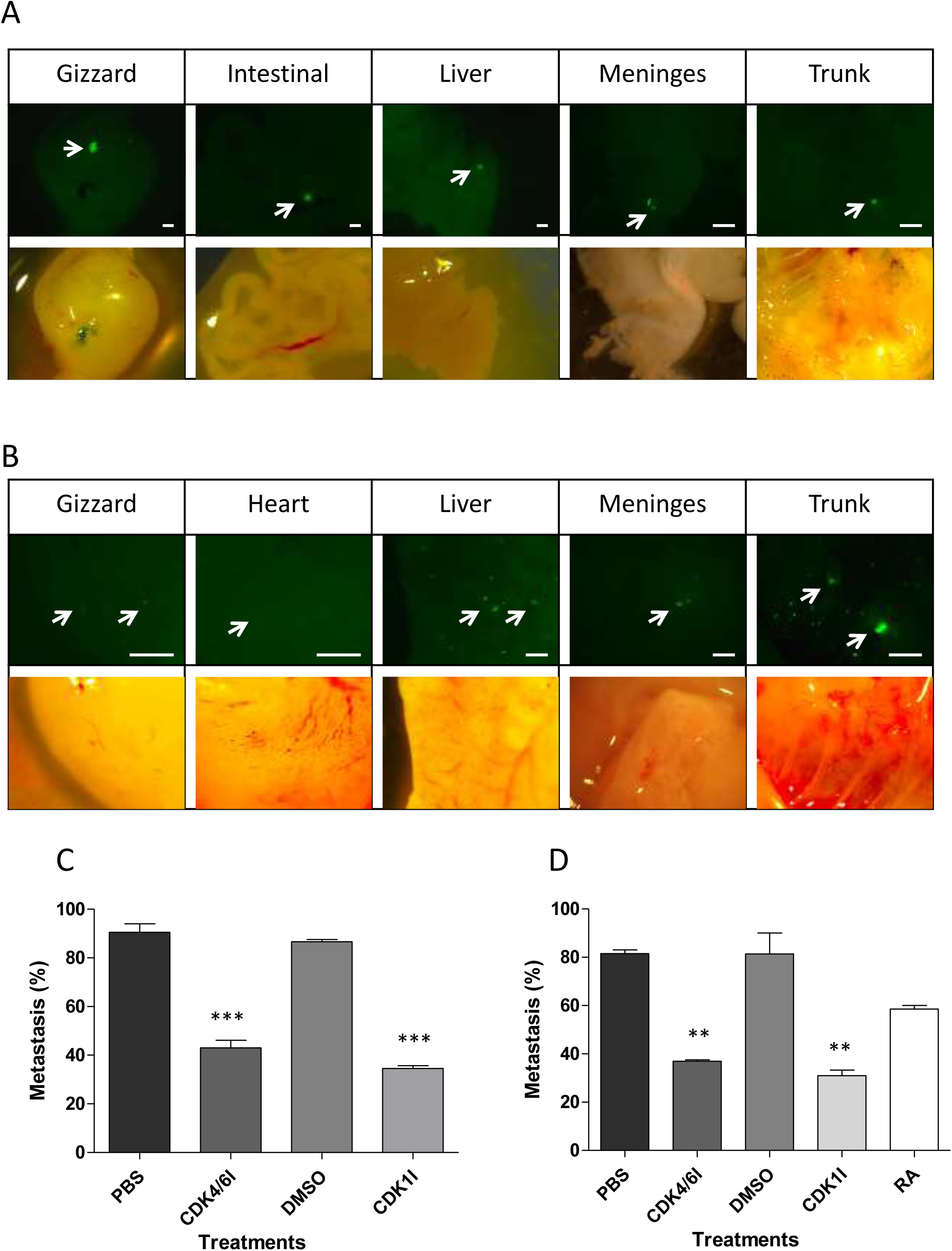
Hypoxic pre-conditioning of cells triggers metastasis *in vivo* and CDK inhibitors reduce this. **A** Representative images of metastatic SKNAS cells in embryonic organs. Hypoxic GFP-labelled SKNAS cells were implanted on the CAM of E7 chick embryos. At E14 embryos were dissected using fluorescence and organs containing metastatic cells were imaged. Scale bar is 500 μm. **B** Representative images of metastatic BE(2)C cells as for **A**. **C** Embryos with SK-N-AS cells implanted on the CAM at E7 were treated with a single injection of 200 μl of either 4.5 mM CDK1i or 32.5 DMSO at E10. All embryos with CAM tumours formed from SK-N-AS cells were dissected at E14 and the percentage of embryos with metastatic cells for each treatment was calculated. A minimum of 15 embryos per condition was analysed (n=15-18). **D** Embryos with BE(2)C cells implanted on the CAM at E7 were treated with a single injection of 200 μl of 9 mM ATRA, 4.5 mM CDK4/6i, 4.5 mM CDK1i, 14% DMSO, PBS, 32.5% DMSO as control at E10. All embryos with CAM tumours formed from BE(2)C cells were dissected at E14 and the percentage of embryos with metastatic cells for each treatment was calculated. A minimum of 12 embryos per condition was analysed (n=12-19).

Similar experiments were conducted with BE(2)C cells. The distribution of metastatic cells was similar to that observed for SK-N-AS (Figure 7B) and CDK1i reduced metastasis by almost 60% (from 76% to 31%) while CDK4/6i reduced metastasis by 56% (from 83% to 36%) (Figure 7D). ATRA was introduced into the allantoic sac in 11.5 μl of DMSO (due to the greater solubility of ATRA in DMSO). Its effect with one and two injections was measured as comparison. One injection reduced metastasis from 80% to 60% while two injections reduced it from 64% (two injections of DMSO) to 27%. Thus ATRA was less efficient at reducing metastasis than either of the CDK inhibitors but it was interesting that the second injection was able to significantly reduce metastasis supporting the notion that metastasis was mainly occurring between E12 and E14 in this model (Figure 7D).

### CDK1i partially reverses hypoxia driven changes in gene expression

We have previously observed significant changes in the expression of more than 20 genes thought to be important in the metastatic process in response to both hypoxia and DMOG pre-conditioning [13]. In order to provide some insight into the mechanism by which the CDK inhibitors might be acting on the metastatic process, we selected eight of these genes to test for their response to CDK1i (Figure 8). Tumours were formed from normoxic cells or from cells treated with DMOG (SK-N-AS) or hypoxia (BE(2)C). The tumours formed by hypoxic preconditioned cells were then treated with either CDK1i or an equivalent volume of DMSO at E10. The mRNA levels in the hypoxia-preconditioned tumours were compared with those in the normoxic tumours. For both BE(2)C and SK-N-AS tumours, the eight genes responded as reported previously [13] with almost all changes reaching significance (Figure 8A&B).

**Figure 8.**
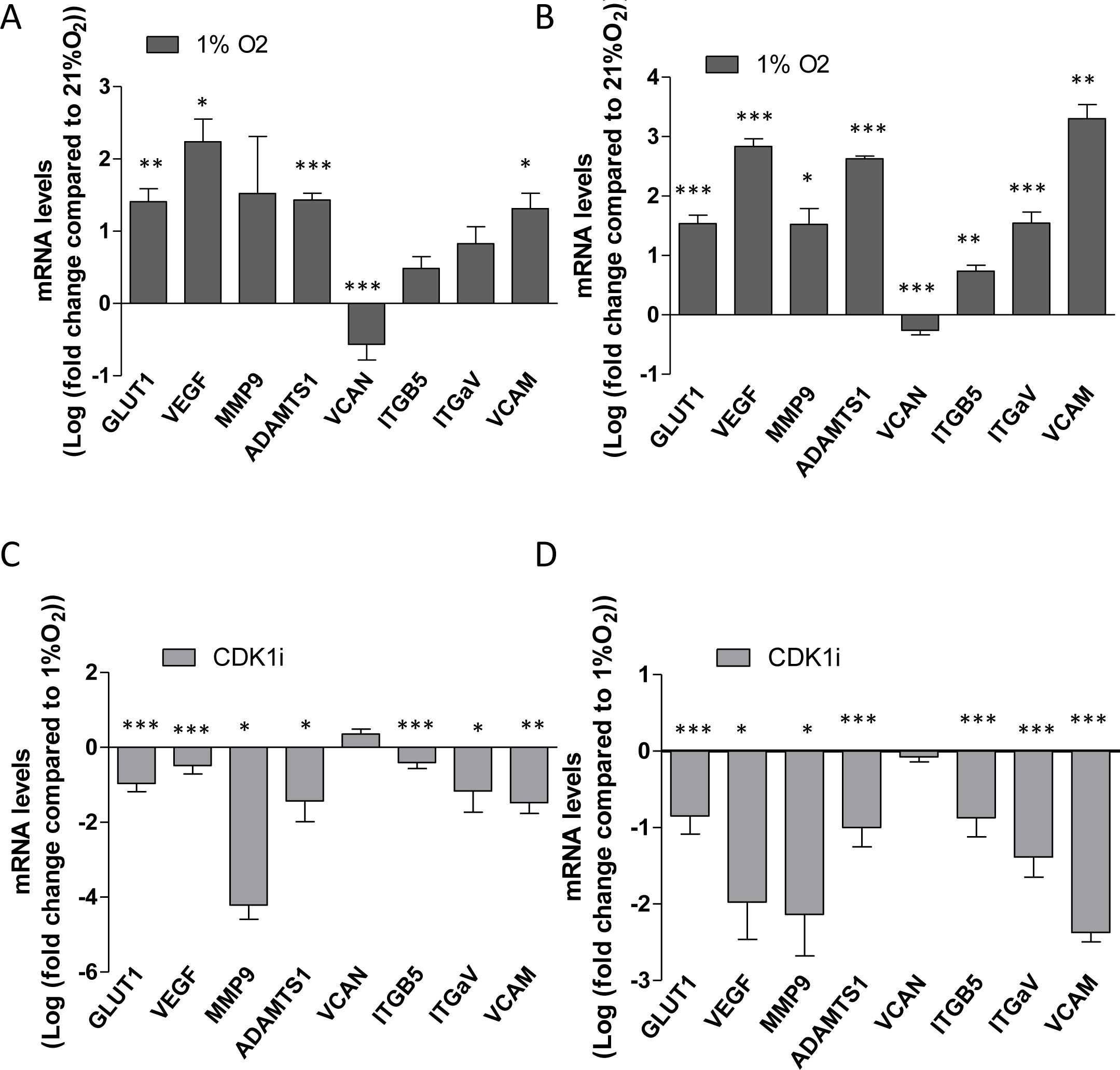
Hypoxia-triggered changes in gene expression is reversed by CDK1i in both BE(2)C and SKNAS tumours. **A** Comparison of gene expression between BE(2)C tumours formed from cells grown in 21% O_2_ and cells pre-cultured in 1% O_2_ for 3 days prior to implantation on the chick CAM at E7. **B** Comparison of gene expression between normoxia and hypoxic SKNAS tumours prepared as for BE(2)C tumours above. **C** Comparison between BE(2)C tumours formed from cells pre-cultured in 1% O_2_ for 3 days prior to implantation on the chick CAM at E7 treated at E10 with either CDK1i or DMSO. **D** Comparison between SK-N-AS tumours formed from cells pre-cultured in 1% O_2_ for 3 days prior to implantation on the chick CAM at E7 treated at E10 with either CDK1i or DMSO. At least one tumour for each condition was obtained from a single experiment and used for comparison. mRNA levels were normalised to GAPDH, UBC and HPRT1 in each case and each bar in the graph represents the normalised mean ±SEM of at least three independent experiments.*P≤0.05, **P≤0.01 and ***P≤0.001 compared with normoxia (A and B) or hypoxia (C and D).

A comparison of hypoxia-treated tumours with and without CDK1i revealed that CDK1i reversed the observed upregulation of expression in the seven up-regulated genes whilst VCAN, whose expression was down-regulated showed a small increase in expression (Figure 8C&D). In some cases expression levels returned to those seen in the normoxic tumours. Taken together these result strongly suggest that a single dose of CDK1i reduces the metastatic phenotype and activity of cells pre-conditioned in hypoxia/treated by DMOG by reversing the increase in expression of genes that are implicated in hypoxia-driven metastasis. In conclusion, we have demonstrated the efficacy of CDK inhibitors in inhibiting neuroblastoma growth and metastasis *in vivo*, regardless of the MYC-N amplification status of the cells, thereby offering potential alternative therapeutic avenue compared to the commonly used ATRA.

## Discussion

The chick embryo is an increasingly valuable model for analysing tumour formation, angiogenesis and most recently the metastatic behaviour of tumour cells. Development of new and effective therapies require a suitable and efficient preclinical model. Here we have demonstrated the value of the chick embryo model for screening potential therapeutic agents for their ability to reduce tumour growth and limit metastasis of neuroblastoma cell lines. CDK inhibitors reduced cell proliferation in tumours in contrast to their *in vitro* activity of driving high levels of cell death. The conspicuous difference between *in vitro* and *in vivo* activity may be due to differing concentration of inhibitor experienced by the cells however lower concentrations in culture did not show any change in cell proliferation. The cellular response to agents in the complex 3D environment of a tumour can be different to that seen in 2D cultures as has been observed by others [17, 18] and this may be the more likely explanation for the differences observed demonstrating yet again the value of the *in vivo* model. Metastasis was judged to have occurred if fluorescent cells were found in the embryo by dissection under a fluorescent stereomicroscope. Single cells within in the embryo could be identified making this more efficient than previous methods of detecting metastatic cells [19]. Careful dissection of many embryos here and previously [13, 14] revealed the typical locations for the metastatic cells. The most common locations were associated with the intestines in the peritoneal cavity or the liver. No metastatic cells were seen prior to the formation of a primary tumour on the CAM and they were only seen in embryos that formed a primary tumour. This confirms that the cells within the embryo were derived from the primary tumour rather than from the initial implantation of cells on the CAM at E7 [13, 14, 20-23]. Thus the CAM tumour model is excellent for investigating response mechanisms to new therapeutic agents in a cost-effective and timely manner.

CDK1:cyclin B complex is sufficient to drive mammalian cells through the cell cycle via phosphorylation of Rb and the consequent activation of the E2F expression program [24]. Inhibiting CDK1 generally arrests the cell cycle at the G2/M boundary and can result in cells undergoing apoptosis [25]. Thus CDK1 makes an attractive target although it should be noted that CDK1 is also responsible for a key phosphorylation step that primes MYCN for degradation [26]. CDK4/6 promotes progression through the G1/S boundary also by phosphorylation of Rb. Components of the CDK4/6 pathway are often dysregulated in cancer making this complex a promising target for treatment [27]. Like CDK1, CDK4 and 6 activate a number of additional pathways and the consequence of inhibiting these kinase may be variable in different tumours [27]. The Inhibitor palbociclib has variable sensitivity across a number of neuroblastoma cell lines [28], furthermore, LEE011, an alternative CDK4/6 inhibitor prompted apoptosis in the majority of neuroblastoma cell lines tested and a reduction in proliferation in BE(2)C xenograft tumours in mice [29] consistent with our results.

Hypoxia drives an aggressive, invasive phenotype and many effects are mediated by HIF1-α. Targets of HIF1-α include proteins involved in invasion, intravasation, adhesion and extravasation and these are likely to be responsible for the change in the neuroblastoma cells to a metastatic phenotype. CDK1, 4 and 6 have been shown to directly increase the stability of HIF1-α by two different mechanisms. CDK1 phosphorylates HIF1-α on Ser688, a modification that reduces the degradation of HIF1-α [30]. CDK4/6 phosphorylates prolyl hydroxylase 1 (PHD1), one of the enzymes responsible for degrading HIF1-α in the presence of O_2_ [31]. Phosphorylation alters the interaction between PHD and HIF1-α potentially stabilising HIF1-α. Both these CDK activities occur independently of O_2_ levels. Inhibiting these CDKs is expected to reduce the abundance of HIF1-α, an effect that should be most obvious under hypoxic conditions when HIF1-α is normally stabilised. Here we find that treatment with the CDK1i, RO-3306, can partially reverse the hypoxia-induced expression pattern in the tumour several days after this pattern has been established in culture.

Blocking the cell cycle via CDK inhibitors is an attractive approach to treating many cancers and had the added value here of potentially prompting differentiation. Some neuroblastoma cell lines and some tumours respond well to retinoic acid by adopting a less aggressive more differentiated phenotype and it is well established that tumours with differentiation markers such as TrkA have a better prognosis. SK-N-AS, along with a number of other cell lines, has not been shown to differentiate so far. However, since both CDK inhibitors reduced cell proliferation in the SK-N-AS CAM tumours, a prerequisite to initiating differentiation, it would be interesting to investigate whether the cells did begin to move down the differentiation pathway *in vivo.*

In conclusion, CDKi provide promising alternatives to ATRA for both reduction of tumour development but also of its dissemination. In addition, they have the potential to be effective in a range of neuroblastoma genetic background, and unlike ATRA, are not restricted to MYCN amplified tumours. The fact that they have been trialled in clinic for other tumour types should facilitate their deployment for neuroblastoma in a near future.

## Methods and Materials

### Cell culture

BE(2)C (human NB, ECACC No. 95011817) and SK-N-AS (human NB, ECACC 94092302) were grown in minimal essential medium DMEM (Life Technologies), 10% Foetal Bovine Serum FBS (Biosera, East Sussex, UK), 100 U/ml penicillin,100 μg/ml streptomycin (Sigma, P0781) and 1% Non-Essential Amino Acids (Sigma, M7145). They were maintained at 37°C with 5% CO_2_ were passaged using 0.05% Trypsin/EDTA (Sigma Aldrich). Cell lines were transduced with green fluorescent protein (GFP) lentivirus as described previously [13, 22]. For hypoxic studies, cells were maintained at 37 °C with 5% CO_2_ and 1% O_2_ (Don Whitley Scientific, Shipley, UK; Hypoxystation-H35) for 72 h. For dimethyloxallylglycine (DMOG) treatment, cells were cultured in media supplemented with 0.5 mM DMOG (Enzo Laboratories, Farmingdale, NY, USA) for 24 h. Treatments were performed with all trans retinoic acid (ATRA, Sigma), the CDK4/6 inhibitor Palbociclib (Selleckchem) or CDK1 inhibitor RO-3306 (Sigma) at indicated concentrations and timings.

### Cell Proliferation Assays and Morphology Analysis

25×10^3^ cells/ml of BE(2)C cells were plated onto coverslips in a 24 well plate, and incubated for 18-24 h to adhere. Cells were treated with DMOG for 24 h then the media was replaced by media containing all trans retinoic acid (ATRA) (10 μM) or Palbociclib (5 μM) or both for a further 72 h. Controls were DMSO (0.06% v/v) or PBS respectively. To assess the morphology of cells, images of cells were obtained using an inverted microscope (Leica DMIRB) prior to fixation. For immunocytochemistry, cells were fixed with 4% paraformaldehyde for 10 min, blocked with 1% BSA,0.1% Triton X100 in 0.12 M phosphate pH7.4 for 30 minutes and stained overnight at 4°C with 1:50 dilution of Ki67 (Abcam ab16667) diluted in blocking buffer. Coverslips were incubated in goat anti rabbit Alexa 594 (Life Technologies) 1:500 for 1 h at room temperature in blocking buffer. Cell nuclei were stained with DAPI (Sigma). Proliferating cells were quantified by Ki67 staining and normalised to the total number of nuclei stained by DAPI. At least three fields per coverslip and 3 coverslips per experiment were counted and a minimum of 300 cells per condition.

### MTT assay

The MTT assay was used to measure cell viability after Palbociclib and CDK1i treatment. BE(2)C cells and SK-N-AS cells were seeded at 5×10^4^ cells/ml into 96 well plates (Costar, Cat no. 3596) with 100 μl of medium and left to adhere for 18-24 h at 37°C 5% CO_2_. Cells were cultured in media supplemented with or without 0.5 mM DMOG for 24 h. Cells were then treated for a further 72 h with medium containing Palbociclib or CDK1i. To analyse cell viability, 5 μl of (5 mg/ml) MTT (Sigma) was then added to each well (100 μl), for 4 h at 37 °C. 100 μl of stop solution (10% SDS in 0.01 M HCl) was added for 1 min. The plate was incubated at 37°C and 5% CO_2_ overnight and absorption at 570 nm was read. All MTT assays were repeated three times.

### Apoptosis determination in unfixed cells

25×10^3^cells/ml of BE(2)C and SK-N-AS cells were plated onto coverslips in a 24 well plate, incubated for 18-24 h, treated for 72 h with medium containing Palbociclib (5 μM) or CDK1i (5 μM); or PBS/DMSO as control. Apoptotic cells were identified using Apoptotic/Necrotic/Healthy Cells Detection Kit from Promokine (PK-CA707-30018) according to the manufacturer’s instructions. Cells were washed twice with 1X Binding Buffer and then incubated with a staining solution (5 μl of FITC-Annexin V, 5 μl of Ethidium Homodimer III and 5 μl of Hoechst 33342 in 100 μl 1X Binding Buffer, 1:20 dilution) for 15 minutes at RT, protected from light. Apoptotic cells were quantified by FITC-Annexin V staining and normalised to the total number of nuclei stained by Hoechst 33342.

### Chick embryo CAM Assays

Fertilised white leghorn chicken eggs were obtained from Tom Barron, Preston, UK. Eggs were incubated at 38°C and 35-40% humidity and windowed at E3 as described previously [22]. For tumour formation 2×10^6^ GFP-labelled cells were applied to the CAM of each embryo [13]. To increase the tumour yield, the CAM was traumatised using a strip of sterile lens tissue causing a small bleed [32] followed by the addition of 5 μl of 0.05% trypsin 0.5mM EDTA. Eggs were resealed and incubated until E14 [23]. Ethical approval for all experiments involving chick embryos was obtained from the Liverpool Animal Welfare and Ethical Review Body.

### Drug delivery

11.3 μl of 0.16 M ATRA in DMSO was diluted to 200 μl with PBS and the colloidal solution was injected into the allantoic sac where the ATRA re-dissolved. ATRA was injected at E11 and E13 for normoxic tumours or E10 followed, in some experiments, by a second injection at E12 for hypoxic tumours. 90 μl of 10 mM Palbociclib in PBS was diluted to 200 μl in PBS and the solution was injected as for ATRA. 65 μl of 14 mM CDK1i in DMSO was diluted to 200 μl with PBS and the solution was injected. Final concentration of the drugs in the egg, assuming 45 ml per egg was 40 μM for RA and 20 μM for the CDK inhibitors per injection. 200 μl of PBS was injected as a control for Palbociclib, 200 μl of 6.6% DMSO as control for ATRA and 200 μl of 32.5% DMSO as control for CDK1i. Embryos were dissected on E14 and tumours analysed.

### Quantitative PCR

*In vitro* samples: cells were seeded at a density of 2×10^6^ per 75 cm^2^ flask and after 24 h, treated with either ATRA (10 μM), Palbociclib (5 μM) combination of both or DMSO alone as control. Every 48 h the medium was replaced with fresh medium as appropriate. After 3 d, RNA was extracted using RNA mini Kit (QIAGEN) according to manufacturer’s instructions. qPCR was carried out on CFX Connect (Biorad) thermocycler using iTaq Universal SYBR green mix (Biorad) with 0.5 μM primers and up to 2 μl cDNA for 35 cycles. An annealing temperature of 60°C was used for all primer pairs and three technical replicates and at least three biological replicates were carried out for each sample. qPCR data analysis was carried out using Bio-Rad CFX Manager 3.0 software. Normalised relative expression of target genes compared to housekeeping genes (GAPDH, HPRT1 and UBC) was calculated using the ΔΔCq analysis mode [33].

*In vivo*: tumours formed from normoxic cells, hypoxic cells and hypoxic cells after CDK1i application at E10 (20 μM) were harvested from the CAM, rinsed in phosphate-buffered saline (PBS), then transferred into RNAlater solution (QIAGEN), and stored at initially at 4°C or −20°C for longer term storage prior to RNA extraction. Tissue was first removed from the RNAlater and transferred to a clean RNase free falcon tube. Liquid nitrogen was used to freeze the tissue before a pestle and mortar was used to disrupt it. RNA was then extracted using RNA mini Kit (QIAGEN). qPCR was performed as described above. A list of the primers used is provided in Table 1.

**Table 1:**
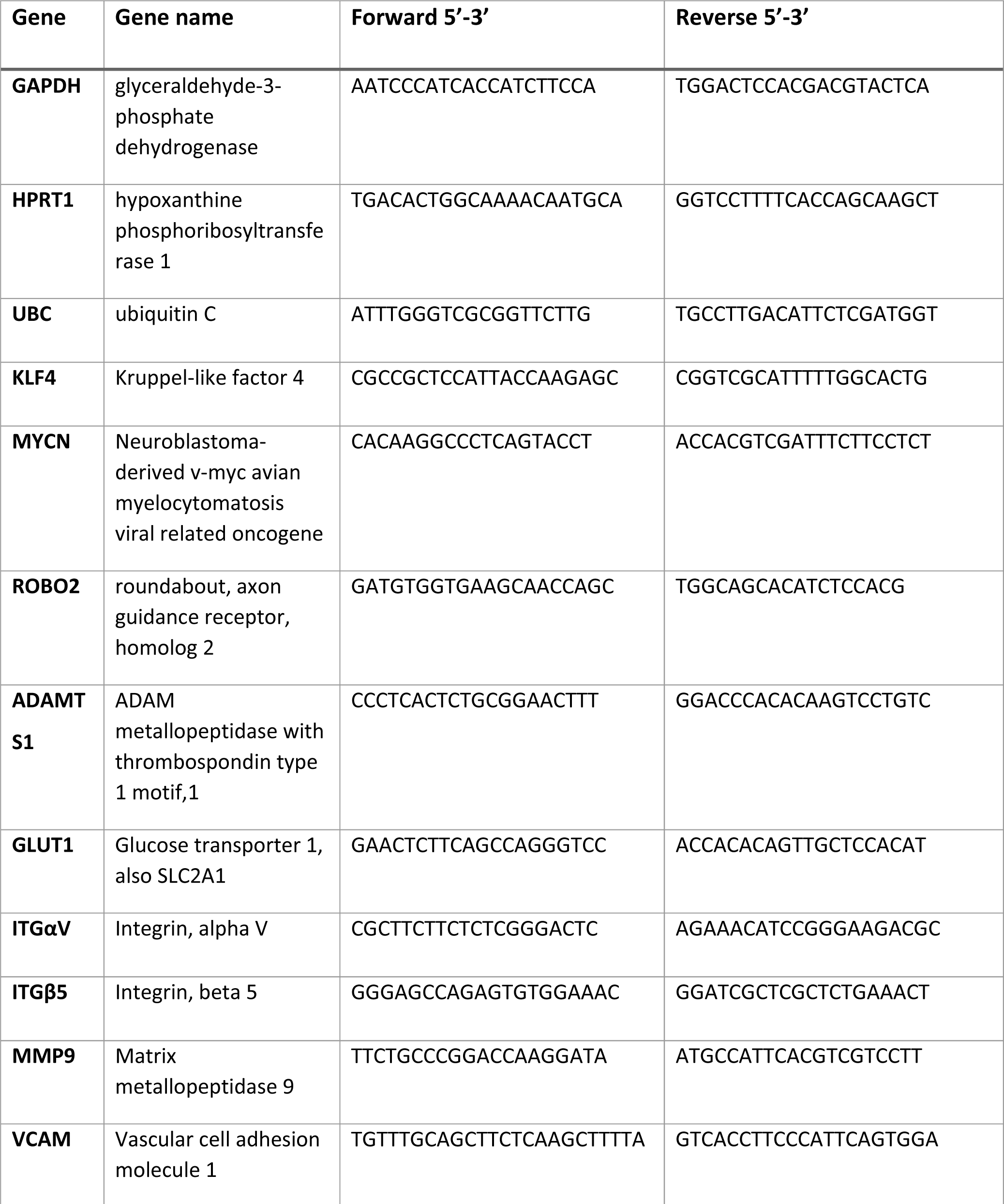
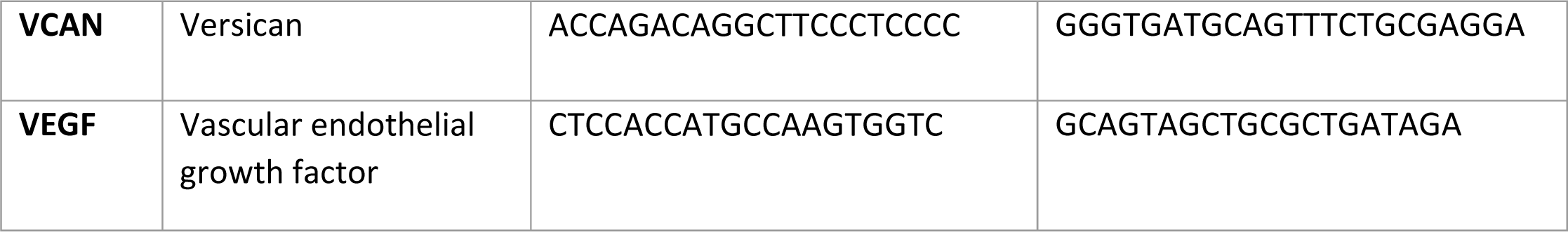
List of primer names and sequences for qPCR

One tumour for each condition (normoxia, hypoxia, hypoxia followed by injection of 200 μl of 4.5 mM CDK1i at E10 or hypoxia followed by 200 μl of 32.5% DMSO at E10) was obtained from a single experiment and fold change was analysed. The experiment was repeated three times and statistical analysis performed on the fold change derived from each experiment.

### Immunohistochemistry

Tumours harvested for paraffin embedding were fixed for 2 h in 4% paraformaldehyde (PFA) or 10% formalin and then embedded in paraffin using standard protocols. 4 μm sections underwent antigen retrieval and were stained using the DAKO Autostainer (K8012) as described previously [14]. Sections were incubated for 30 min with Ki67 antibody (1:200) (DAKO M7240) in 5% BSA in Tris Buffered Saline followed by goat anti-mouse HRP (Abcam)(1:250) and staining with 3,3’-diaminobenzidine DAB (1 drop per ml). Haematoxylin staining was performed on all the slides to assist in distinguishing between tumour and chick nuclei. A total of 9 fields from 3 slides were counted per tumour and at least three tumours per condition were analysed.

### TUNEL Assays

TUNEL assay was performed on tumour sections prepared as above using the DeadEnd™ Colorimetric TUNEL System (G7130) kit according manufacturer’s instructions (Promega, Madison, WI, USA). TUNEL-positive nucleus was identified as apoptotic cells. Cell counting was performed on at least three tumours per condition, 9 fields from 3 slides per tumour were counted and analysed.

### Statistical analysis

Statistical significance was computed using Student’s t-test or one-way ANOVA followed by a post hoc tukey test using SPSS. All data are presented as mean ±SEM. (standard error of the mean).

### Data Availability

The datasets generated during and/or analysed during the current study are available from the corresponding author on reasonable request.

## Acknowledgements

The authors thank Dr Lakis Liloglou for assistance with the qPCR analysis, Dr Rajeev Shukla for assistance with the histology, Dr Helen Kalirai for assistance with the Ki67 staining and Professor Barry Pizer and Dr Rabiu Inuwa for helpful discussions. This work was supported by grants from the Iraqi Higher Education Ministry, Iraqi cultural attaché London (RS), The Neuroblastoma Society UK and NC3Rs (AH) and The Royal College of Surgeons and NWCRF (KS).

## Author contributions

RS designed and carried out the majority of the experiments, prepared figures and tables, and wrote parts of the manuscript, KS performed experiments, analysed data and prepared figures, AH performed experiments, analysed data and reviewed the manuscript. PL reviewed the data and provided clinical input throughout the project, VS was responsible for obtaining funding, supervision of the project and for writing the manuscript. DM was responsible for the original design and conception of the project obtaining funding, supervision of the project and for writing the manuscript.

## References

1. Cheung, N.K. and M.A. Dyer, Neuroblastoma: developmental biology, cancer genomics and immunotherapy. Nat Rev Cancer, 2013. 13(6): p. 397–411.

2. Thiele, C.J., C.P. Reynolds, and M.A. Israel, Decreased expression of N-myc precedes retinoic acid-induced morphological differentiation of human neuroblastoma. Nature, 1985. 313(6001): p. 404–6.

3. van Noesel, M.M., et al., Neuroblastoma 4S: a heterogeneous disease with variable risk factors and treatment strategies. Cancer, 1997. 80(5): p. 834–43.

4. Caren, H., et al., High-risk neuroblastoma tumors with 11q-deletion display a poor prognostic, chromosome instability phenotype with later onset. Proc Natl Acad Sci U S A, 2010. 107(9): p. 4323–8.

5. Edsjo, A., et al., Neuroblastoma cells with overexpressed MYCN retain their capacity to undergo neuronal differentiation. Lab Invest, 2004. 84(4): p. 406–17.

6. Guglielmi, L., et al., MYCN gene expression is required for the onset of the differentiation programme in neuroblastoma cells. Cell Death Dis, 2014. 5: p. e1081.

7. Borriello, A., et al., p27Kip1 accumulation is associated with retinoic-induced neuroblastoma differentiation: evidence of a decreased proteasome-dependent degradation. Oncogene, 2000. 19(1): p. 51–60.

8. Hardwick, L.J. and A. Philpott, Nervous decision-making: to divide or differentiate. Trends Genet, 2014. 30(6): p. 254–61.

9. Palanisamy, R.P., Palbociclib: A new hope in the treatment of breast cancer. J Cancer Res Ther, 2016. 12(4): p. 1220–1223.

10. Goel, S., et al., CDK4/6 Inhibition in Cancer: Beyond Cell Cycle Arrest. Trends Cell Biol, 2018.

11. Kang, J., et al., Targeting cyclin-dependent kinase 1 (CDK1) but not CDK4/6 or CDK2 is selectively lethal to MYC-dependent human breast cancer cells. BMC Cancer, 2014. 14: p. 32.

12. Xia, Q., et al., The CDK1 inhibitor RO3306 improves the response of BRCA-proficient breast cancer cells to PARP inhibition. Int J Oncol, 2014. 44(3): p. 735–44.

13. Herrmann, A., et al., Cellular memory of hypoxia elicits neuroblastoma metastasis and enables invasion by non-aggressive neighbouring cells. Oncogenesis, 2015. 4: p. e138.

14. Swadi, R., et al., Optimising the chick chorioallantoic membrane xenograft model of neuroblastoma for drug delivery. BMC Cancer, 2018. 18(1): p. 28.

15. Clark, O., S. Daga, and A.W. Stoker, Tyrosine phosphatase inhibitors combined with retinoic acid can enhance differentiation of neuroblastoma cells and trigger ERK- and AKT-dependent, p53-independent senescence. Cancer Lett, 2013. 328(1): p. 44–54.

16. Wyatt, R.D. and B. Howarth, Jr., Effect of dimethyl sulfoxide on embryonic survival and subsequent chick performance. Poult Sci, 1976. 55(2): p. 579–82.

17. Anastasiou, D., Tumour microenvironment factors shaping the cancer metabolism landscape. Br J Cancer, 2017. 116(3): p. 277–286.

18. Gandellini, P., et al., Complexity in the tumour microenvironment: Cancer associated fibroblast gene expression patterns identify both common and unique features of tumour-stroma crosstalk across cancer types. Semin Cancer Biol, 2015. 35: p. 96–106.

19. Zhao, S.G., et al., Development and validation of a novel platform-independent metastasis signature in human breast cancer. PLoS One, 2015. 10(5): p. e0126631.

20. Lokman, N.A., et al., Chick Chorioallantoic Membrane (CAM) Assay as an In Vivo Model to Study the Effect of Newly Identified Molecules on Ovarian Cancer Invasion and Metastasis. Int J Mol Sci, 2012. 13(8): p. 9959–70.

21. Ribatti, D., The chick embryo chorioallantoic membrane in the study of tumor angiogenesis. Rom J Morphol Embryol, 2008. 49(2): p. 131–5.

22. Carter, R., et al., Exploitation of chick embryo environments to reprogram MYCN-amplified neuroblastoma cells to a benign phenotype, lacking detectable MYCN expression. Oncogenesis, 2012. 1: p. e24.

23. Herrmann, A., D. Moss, and V. See, The Chorioallantoic Membrane of the Chick Embryo to Assess Tumor Formation and Metastasis. Tumor Angiogenesis Assays: Methods and Protocols, 2016. 1464: p. 97–105.

24. Santamaria, D., et al., Cdk1 is sufficient to drive the mammalian cell cycle. Nature, 2007. 448(7155): p. 811–5.

25. Vassilev, L.T., et al., Selective small-molecule inhibitor reveals critical mitotic functions of human CDK1. Proc Natl Acad Sci U S A, 2006. 103(28): p. 10660–5.

26. Sjostrom, S.K., et al., The Cdk1 complex plays a prime role in regulating N-myc phosphorylation and turnover in neural precursors. Dev Cell, 2005. 9(3): p. 327–38.

27. Tigan, A.S., et al., CDK6-a review of the past and a glimpse into the future: from cell-cycle control to transcriptional regulation. Oncogene, 2016. 35(24): p. 3083–91.

28. Rihani, A., et al., Inhibition of CDK4/6 as a novel therapeutic option for neuroblastoma. Cancer Cell Int, 2015. 15: p. 76.

29. Rader, J., et al., Dual CDK4/CDK6 inhibition induces cell-cycle arrest and senescence in neuroblastoma. Clin Cancer Res, 2013. 19(22): p. 6173–82.

30. Warfel, N.A., et al., CDK1 stabilizes HIF-1alpha via direct phosphorylation of Ser668 to promote tumor growth. Cell Cycle, 2013. 12(23): p. 3689–701.

31. Ortmann, B., et al., CDK-dependent phosphorylation of PHD1 on serine 130 alters its substrate preference in cells. J Cell Sci, 2016. 129(1): p. 191–205.

32. Armstrong, P.B., J.P. Quigley, and E. Sidebottom, Transepithelial invasion and intramesenchymal infiltration of the chick embryo chorioallantois by tumor cell lines. Cancer Res, 1982. 42(5): p. 1826–37.

33. Schmittgen, T.D. and K.J. Livak, Analyzing real-time PCR data by the comparative C(T) method. Nat Protoc, 2008. 3(6): p. 1101–8.

